# Euler method can outperform more complex ODE solvers in the numerical implementation of the Izhikevich artificial Spiking Neuron Model given the allocated FLOPS

**DOI:** 10.1101/2021.11.30.470474

**Authors:** Giuseppe de Alteriis, Enrico Cataldo, Alberto Mazzoni, Calogero Maria Oddo

## Abstract

The Izhikevich artificial spiking neuron model is among the most employed models in neuromorphic engineering and computational neuroscience, due to the affordable computational effort to discretize it and its biological plausibility. It has been adopted also for applications with limited computational resources in embedded systems. It is important therefore to realize a compromise between error and computational expense to solve numerically the model’s equations. Here we investigate the effects of discretization and we study the solver that realizes the best compromise between accuracy and computational cost, given an available amount of Floating Point Operations per Second (FLOPS). We considered three fixed-step solvers for Ordinary Differential Equations (ODE), commonly used in computational neuroscience: Euler method, the Runge-Kutta 2 method and the Runge-Kutta 4 method. To quantify the error produced by the solvers, we used the Victor Purpura spike train Distance from an ideal solution of the ODE. Counterintuitively, we found that simple methods such as Euler and Runge Kutta 2 can outperform more complex ones (i.e. Runge Kutta 4) in the numerical solution of the Izhikevich model if the same FLOPS are allocated in the comparison. Moreover, we quantified the neuron rest time (with input under threshold resulting in no output spikes) necessary for the numerical solution to converge to the ideal solution and therefore to cancel the error accumulated during the spike train; in this analysis we found that the required rest time is independent from the firing rate and the spike train duration. Our results can generalize in a straightforward manner to other spiking neuron models and provide a systematic analysis of fixed step neural ODE solvers towards an accuracy-computational cost tradeoff.

## 1 Introduction

### 1.1 Motivation

Since many years, Izhikevich’s Artificial Spiking Neuron Model (ISNM) has been often adopted to simulate various types of neurons (Izhikevich [2003, 2004]). It allows a compromise between computational cost and the abundance of neuronal features it is able to exhibit (by changing only four parameters it can model a broad range of dynamics) (Izhikevich [2004]). This model has been widely employed in neuromorphic computing, machine learning and neural engineering both for a neuromorphic encoding of signals (Oddo et al. [2016], Osborn et al. [2018], Mazzoni et al. [2020]) or to reproduce neuronal ensembles for machine learning (Li et al. [2017], Rice et al. [2009], Rongala et al. [2017]) and neuroscience tasks (Shafiei et al. [2019], Vinaya and Ignatius [2018]). The ISNM does not mimic the time evolution of the membrane potential of a neuron at a physiological level, but just at a functional one. This allows a simplification and allows a good real-time implementation in microprocessors or FPGAs. However, this poses a big challenge in robotics and biomedical engineering because the amount of real-time operations that can be performed is limited by the utilization of embedded systems. To allocate optimally the resources of microprocessors, we seek a trade-off between number of operations performed and accuracy, by investigating the best Ordinary Differential Equation (ODE) solver and discretization time *τ* (Heidarpur et al. [2020]). *In fact, the model is sensitive to the τ* used (Heidarpur et al. [2020]), and the bigger it gets, the more the spike timing accumulates a delay with respect to the ideal solution. This effect is evident in the three ODE solvers we are going to show in the paper.

### 1.2 State of the Art and Related Work

Some values utilized for *τ* in published papers are dt=0.01ms Yu et al. [2017a], dt=0.1ms Yu et al. [2017b] and dt=0.5ms (Izhikevich, 2003; Spigler et al., 2012). Our previous works (Gunasekaran et al. [2019], de Alteriis and Oddo [2021]) analyze the effect of varying *τ* on the convergence of the ISNM with Euler’s method. They present a way to find a compromise between the number of operations and the precision of spike sequences generated.(Humphries and Gurney [2007]) propose a Zero-Order Hold solution of the model, which consists, at each step, of solving each equation in an exact manner by keeping the other variable constant. As a quality metric of the solver, they consider the mean error per spike (time difference from the spike of an ideal solution generated with a higher order variable step solver). (Long and Fang [2010]) show the discretization step causal role on the firing rate in different neural models, and (Skocik and Long [2014]) deepens this by taking in consideration also the deviation of the voltage profile from the ideal solution. (Hopkins and Furber [2015]) explores different solution methods for the Izhikevich model and shows that using a 1 ms time step the Runge-Kutta 2 method looks to be the best by comparing it with other methods and showing how the lag of the spikes with respect to an reference spike train accumulates. Finally, (Valadez-Godínez et al. [2020]) propose an approach, based on the multi-objective optimization theory (Chang [2015]), to find the optimal simulation parameters.

### 1.3 Contribution

We inverted this perspective: instead of considering the time step as a parameter to choose, we speculated on the total amount of Floating Point Operations per Second available for the specific application. Given the computational availability (i.e. number of FLOPS) it is crucial to understand whether it is beter to choose a more sophisticated solver with a bigger *τ* or a more basic method with a smaller *τ* (as highlighted in Figure 1). Why is this question relevant? The ISNM model presents a reset rule, which coincides with the spike event. It is therefore legit to ask: how does this reset rule affect the numerical solution? Is it better to make a bigger error in the reproduction of the membrane potential and a smaller error in the time of the reset, or vice-versa? As an example, when 30 kFLOPS are available, Euler’s method or RK2 with a smaller time step outperform RK4 with a bigger one (Figure 1). In most cases, the quantity that plays a relevant role is the spike timing rather than an exact quantification of the membrane potential. Finally, we believe that in some neuromorphic engineering applications it is useful to assess the time needed by the solvers to converge to the equilibrium solution. In fact, (Hopkins Furber, 2015; Long Fang, 2010) show how the error of the solver accumulates at each spike, but it is interesting to investigate how this gets reset.

**Figure 1:**
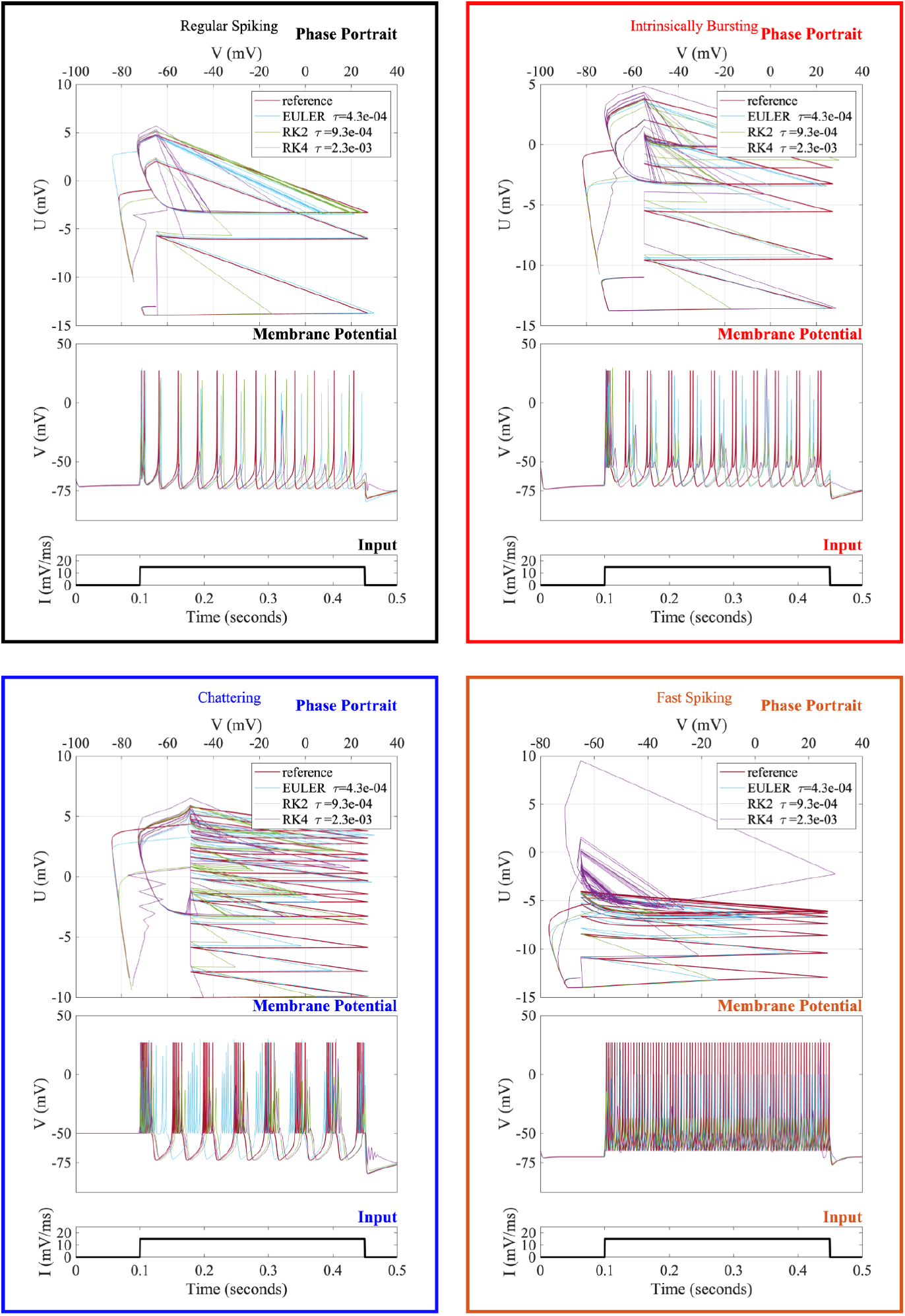
ISNM numerically solved with: Euler method (E, coded in blue), Runge-Kutta 2 (RK2, green), Runge-Kutta 4 (RK4, purple). The reference solution is in red. Given a fixed number of 30 kFLOPS, *τ* was calculated for each solver. In each panel: on the bottom, the input current. On the top, the phase plane plots. In the middle: the neuron’s membrane potential. Each panel represents a different neuron model, RS (black, A), IB (red, B), CH (blue, C), FS (orange, D).

## 2 Materials and Methods

### 2.1 The Izhikevich Artificial Spiking Neuron Model

We investigate the Izhikevich Model (Izhikevich [2003]), consisting of two ordinary differential equations (ODEs):

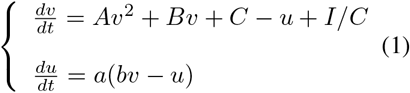

*v* represents the membrane potential and *u* is called recovery variable and plays a role in adaptation by introducing a feedback on *v*. The model presents a reset which is: if *v > v*_*th*_ then *u* = *u* + *d, v* = *c*. The variables A = 0.04 /(ms mV), B = 5 /ms, C = 140 mV/ms are the fixed parameters of the model, while the threshold *v*_*th*_ is set to 30 mV. The input current is *I/C* (mV/ms). These two ODEs have a strong sensitivity to the parameters (*a, b, c, d*). By changing them we can model various types of neuronal characteristics (Izhikevich [2004]): *a* is the *u*’s time scale; *b* is *u*’s sensitivity to fluctuations of *v*; *c* is the reset value of *v* after the spike; d is the value added to *u* after the spike. Here we adopted the four most important and most used parameters sets:

Fast Spiking and Regular Spiking parameters set: While the input is on, the neuron fires continuously. This tonic spiking behavior can be observed in regular spiking (RS) (a = 0.02, b = 0.2, c = 65, d = 8) excitatory neurons, and fast spiking (FS) inhibitory neurons (a = 0.1, b = 0.2, c = 65, d = 2). Firing of RS and FS neurons means that the input is persistent. Regular-spiking cells show a strong adaptation to constant inputs, while fast-spiking neurons have little adaptation. Intrinsically bursting (IB) (a = 0.02, b = 0.2, c = 55, d = 4) and Chattering (CH) cells (a = 0.02, b = 0.2, c = 50, d = 2) generate clusters of spikes (bursts), which can be single or repetitive. Regular Spiking and Intrinsically Bursting cells can be either spiny stellate cells or pyramidal neurons, by which the excitatory cells of the cerebral cortex are constituted. Fast Spiking neurons are sparsely spiny or smooth, non-pyramidal, and likely they are GABAergic inhibitory interneurons (Connors and Gutnick [1990], Gibson et al. [1999]). We set, as initial conditions of the system, *u* = *bv* and *v* = *c*.

### 2.2 Euler and Runge Kutta solvers

We studied three single step numerical ODE solvers (the solution at a time step is computed based just on the solution at the previous one): they are in the group of Runge-Kutta (RK) solvers, which are frequently employed for neural models: Euler method(E), which relies on an approximation of the first order of the derivative and it is also known as the Runge Kutta 1 (RK1) method, RK2, and RK4. If we consider an ODE system of the type *x*^′^ = *f* (*x, t*), for RK1 a single evaluation of *f* is needed, RK2 performs two *f* evaluations, and RK4 performs four evaluations. For instance, E approximates the potential via the update rule: *v*(*t* + *h*) = *v*(*t*) + *h*(*Av*^2^ + *Bv* + *C* − *u* + *I/C*). We kept a fixed discretization step. We counted, accordingly with (Izhikevich, 2004) that this method requires 13 Floating Point Operations per update (FLOP). The same consideration holds true for RK2, which relies on a second order approximation (28 FLOP) and RK4, which relies on a fourth order (68 FLOP). One could then use (Figure 1) a higher-order method with a bigger *τ* or do the opposite, considering that the total amount of FLOPS can be computed by dividing the FLOP for one iteration by the discretization step *τ* (FLOPS = FLOP per iteration/*τ*). Our implementation of the ODEs was done using Simulink (Mathworks, USA). Note that the Runge Kutta 2 method has different variants (Quarteroni et al., 2010), but in our case we have used Heun’s method as it is the most directly understandable and the one implemented in Simulink.

### 2.3 Quality Metrics

To generate the reference train of spikes, we used the RK5 method with a *τ* of 0.5 *µs* because this is a temporal scale which below the typical timing of neuronal dynamics. As a first quality metrics for E, RK2 and RK4 we considered the Victor Purpura Spike Train Distance (VPD) with the reference spike train. The VPD quantifies the alignment of sequences of discrete events (Victor and Purpura [1996], Satuvuori and Kreuz [2018]). It is non-Euclidean, and is operationally computed as the minimal cost necessary to make two spike sequences overlap, using the combination of the steps listed below that minimize the overall cost:

- Removing a spike, which has a cost of 1
- Insertion of a spike, which has a cost of 1
- Spike shift of an amount of time *δt*, which has a cost of *q/*|*δt*|)

*q* is the cost to move a spike and its unit of measurement is *s*^-1^. This distance has been employed also in neuromorpic computing systems, for instance in neuro-robotic tactile applications (Rongala et al. [2017]). To check whether the results were generalizable we considered a second metric, which is the Van Rossum spike train distance (VRD): it convolves the spike sequences with continuous exponential kernels (with a time constant *t*). Then, the Euclidean integral distance is computed between the continuous functions obtained.The parameter *t* has the same role that 1*/q* has in the Victor Purpura Distance. In particular, relatively lower values of *t* (relatively to the inter-spike-interval) enforce this distance to be more sensitive to finer timing differences, whilst higher *t* values make it primarily sensitive to the overall firing rate difference.

### 2.4 Data Collection and Data Analysis

We collected our dataset in the following way: we implemented the four models using Simulink. First, we generated spike trains giving to the model a step input as in Figure 1. We generated the reference spike trains using a time step of 0.5 ms and the Runge Kutta 5 method. We then ran the simulations multiple times for both E, RK2 and RK4 by varying linearly *τ* from 0.1 to 2 ms. We collected both the spike timing arrays and the time course of the variables *u* and *v* of the model. To assess the time of convergence to the equilibrium solution, given the two state variables u and v, we evaluated the squared Euclidean distance in the phase space between the numerical solution and the ideal one. Since we are considering numerical solutions performed with different steps, we subsampled the results to a common reference frame of 1 ms time step. When the neuron goes OFF, this distance converges exponentially to zero. We evaluate the time of convergence as the moment when the distance crosses a threshold, which was set to 0.001. Finally, we followed this procedure to collect the VPDs by varying the number of FLOPS.

1. we select a possible range of FLOPS (from 10 kFLOPS to 7 MFLOPS) and amplitudes of the input (from 10 to 100 mV/ms) Reiterate this:
2. for each flop in the range (varied logarithmically) we compute the discretization time for each solver using the formula *τ* = FLOP-per-iteration /FLOPS
3. we run the solver to get the numerical solution for each amplitude
4. we compute the VPD between the numerical solution and the Reference

We have performed the same data collection strategy by considering, instead of variable amplitude inputs, a sine function of amplitude 30 mV/ms and varying the frequency from 10 to 100 Hz.

## 3 Results

### 3.1 Characterization of Spike Delay

The first problem we addressed is the increase in the error in the spike timing when the discretization step increases. We provided as input to the system the same current step of 15 mV/ms amplitude of Figure 1 and generated different spike trains by using the three solvers with a *τ* that varied linearly (from 100 *µ*s to 2 ms). The raster plots in Figure 2 display that the accumulated delay is greater than zero: increasing *τ*, the E, RK2 and RK4 solvers exhibit a latency respect to the reference spike train that increases. Therefore, with increasing *τ* the approximate solutions exhibit a slower dynamics, that is observable also in the phase-space (Figure 1). The error accumulates in a linear fashion, as also noted also by (Hopkins and Furber [2015], de Alteriis and Oddo [2021])for the RS and IB. To quantify the delay and its linear behavior, for each *τ* the difference between the spike time of the train produced by the solver and the reference spike time was computed, as in (Figure 2). We observed and quantified a linear law between the error and the spike count (but not the timing of the spike): we computed the slopes of the linear regressions and the R squared coefficients (Figure 3). As expected, RK4 accumulates less error and therefore for each discretization step the slope is the lowest, while it is the highest in E. Considering R squared coefficients, we found that this linearity persists under a *τ* threshold of 1 ms. RS and IB neurons were amenable of this analysis, while FS and CH were not. In fact, as recognizable in the raster plots, in FS and CH the solver generates additional spikes, or lacks some spikes, and thus it is impossible to track every single spike and its delay from the ideal one.

**Figure 2:**
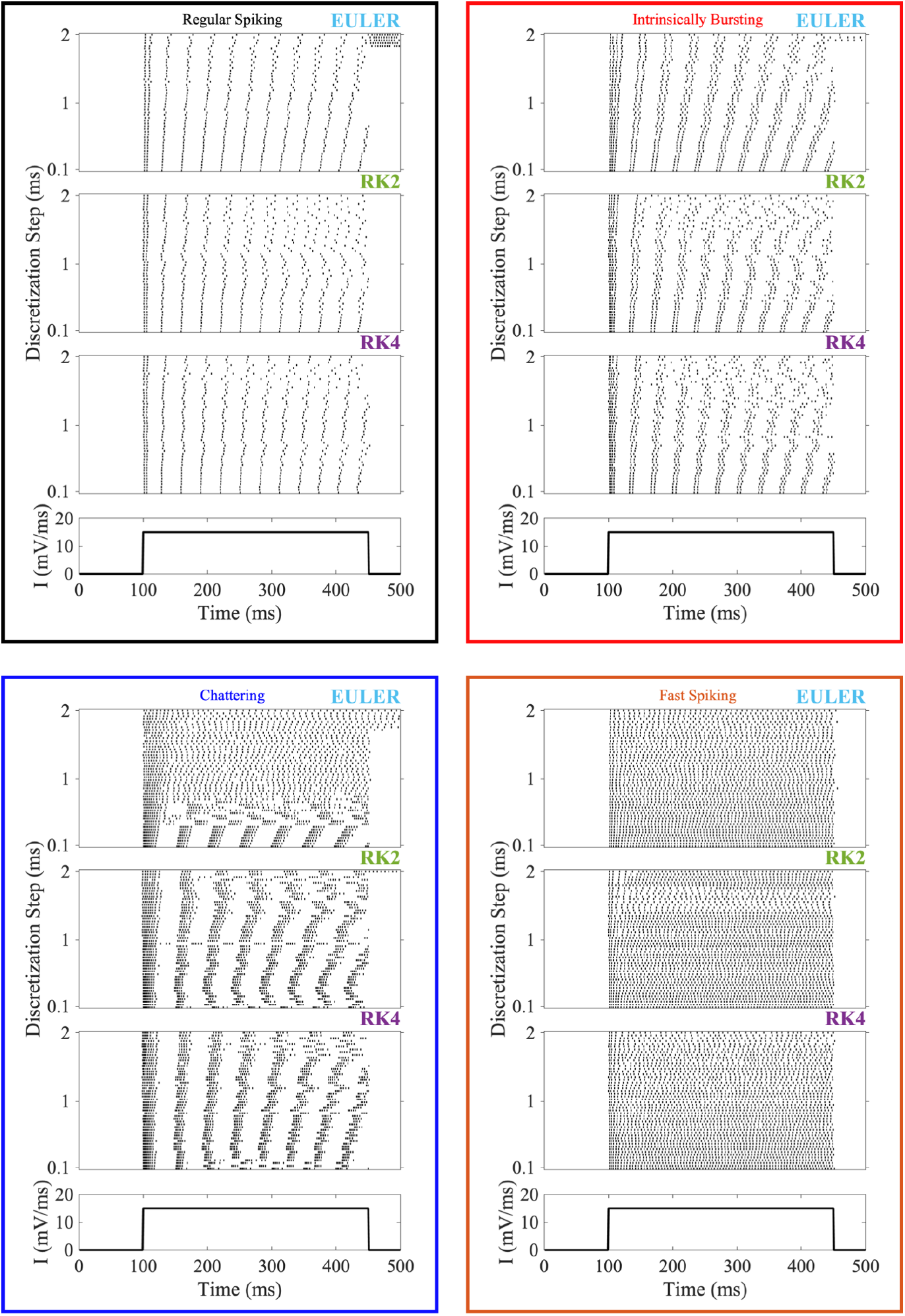
We obtained these raster plots by generating for every solver 50 spike trains, varying *τ* from 0.1 to 2 ms. Each panel represents a different neuron: A, black, regular spiking, B, red, intrinsically bursting, C, blue, chattering, D, orange, fast spiking.

**Figure 3:**
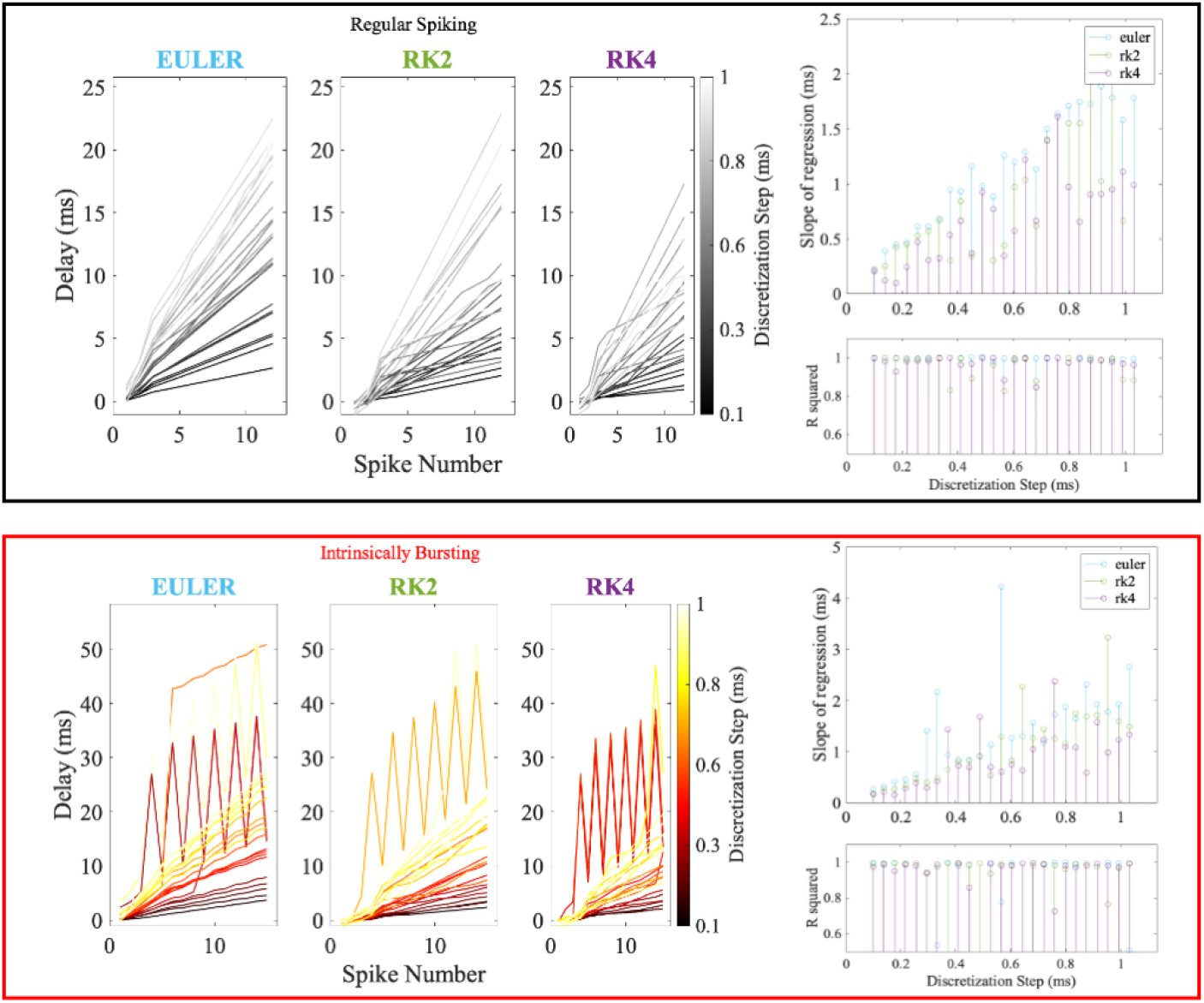
Were generated spike trains, using the current input of (Figure 1) and (Figure 2), and varying *τ* linearly from 0.1 to 1 ms. We repeated this operation for each ODE solver. On the left: each line is the error of the spikes with respect to the reference ones. For better clarity, the lines are interpolated. Their slope depend on *τ*, which is therefore colormap coded. On the right: OLS (Ordinary Least Squares) linear regression of the aforementioned curves. Below there are the R squared coefficients. Above, the slopes, plotted for each method. The same operation is repeated for the RS neuron (panel a, black) and the IB (panel B, red).

The explanation of this linearity is that *dv/dt* increases exponentially during the time course of a spike event. Given the role that the derivatives of the solution have on the error of ODE solvers, this implies that the contribution to the numerical error in the spiking event is bigger than the one accumulated in the phases where the neuron does not fire, making them negligible. Therefore, as also in (Hopkins Furber, 2015) in a first approximation the delay from the reference spike depends linearly on the spike count. The practical value of this result, is to show that the error is proportional to the duration of the spike train. If the input duration and therefore the spike train duration is a-priori known, this could give an estimate of the maximum error. For example, for the Regular Spiking neuron model solved with E in Figure 3, with a discretization step of 1 ms, it is possible to fire uninterruptedly no more than 10 times to keep the error below 20 ms. For example, E with a step of 0.5 ms could be enough to simulate a Slow Adapting Merkel cell with low firing rate for a tactile receptor, whereas for neurons which exhibit higher firing rates it is better to allocate more computational resources.

For RS and IB, confirming (de Alteriis and Oddo [2021]), we observed that the linear law keeps stable only for one uninterrupted spike train. In fact, when the firing of the neuron stops (the neuron goes OFF) the accumulated error will reset. A a new linear accumulation will occur within the following spike train. This is because during the spiking phase the positive delay accumulates, and, as the neuron is re-polarized, gets reset, as showed in Figure 4. If the time spent OFF is not enough, the numerical solution will not be able to converge to the real equilibrium solution, therefore the delay will not reset, as in Figure 4. For a quantification of the decrease in the error and the time necessary to reset it see Section 3B.

**Figure 4:**
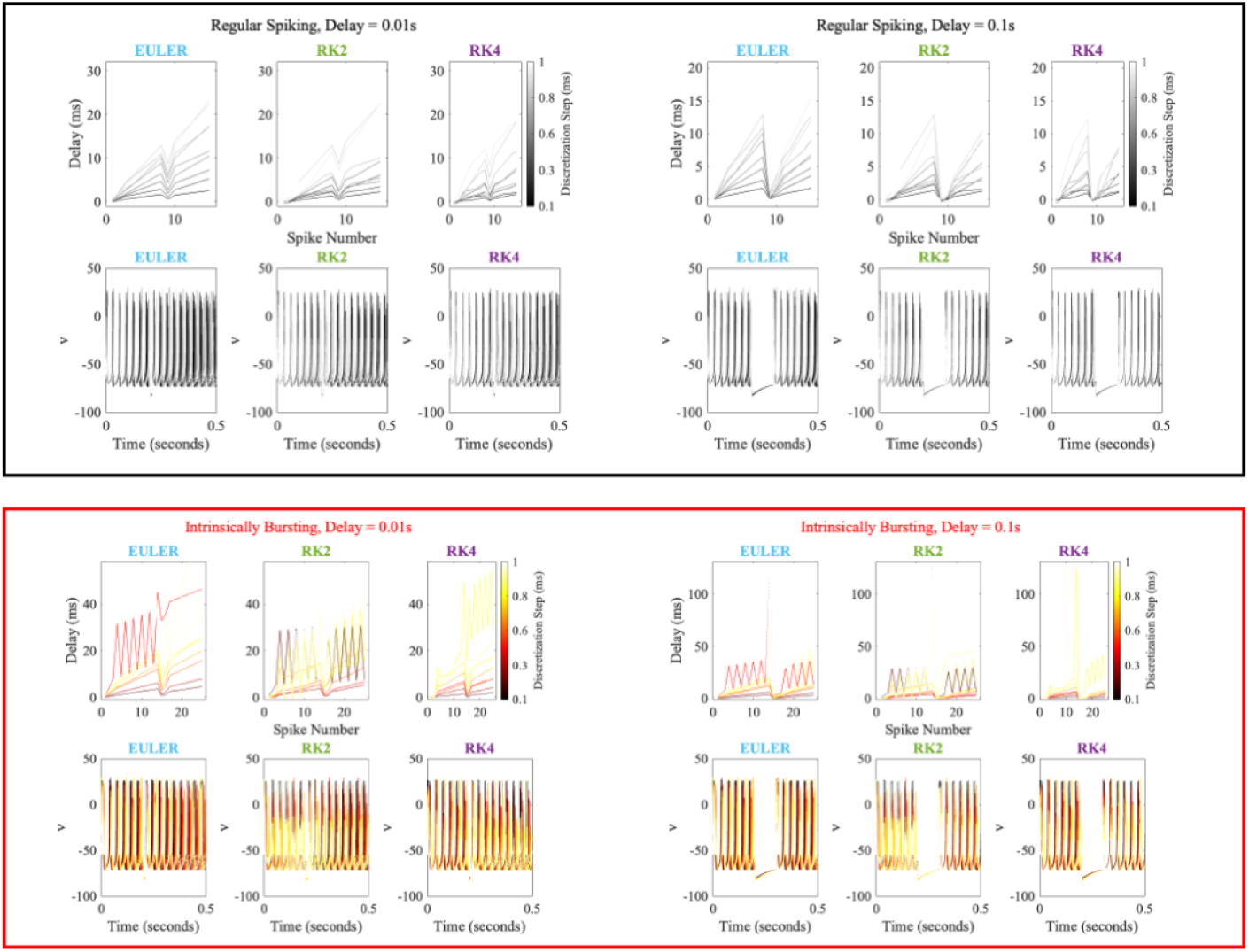
The same analysis of Figure 3, but considering an input that goes to zero after 0.2 seconds and resumes with a delay of 0.01 s, on the left, and 0.1 s, on the right. The same operation is repeated for the RS neuron (panel a, black) and the IB (panel B, red).

### 3.2 Which is the best solver?

We used the Victor Purpura Distance (see Materials and Methods) and the number of FLOPS available as a parameter. Therefore we used the following strategy, as explained in the Materials and Methods section: given a number of available FLOPS, we computed the discretization time relative to that amount of FLOPS for each method (Euler, Runge Kutta2, Runge Kutta 4). This is inspired by engineering scenarios where a dedicated number of FLOPS for a given problem is available, and it is necessary to choose the best solver given the computational restrictions. We iterated this procedure considering current inputs of different amplitudes, which affect the firing rate of the neurons. We then generated 3d surfaces where the x axis is the number of FLOPS, the y axis the amplitude of the current input and the z axis is the VPD, which is our quality metric. This means that we are seeking for the solver whose surface is under the others, as in Figure 5. In all the figures there an area where Euler outperforms Runge Kutta 2 and Runge Kutta 4 and a second area where Runge Kutta2 outperforms both Euler and Runge Kutta 4. To quantify these two areas, let’s call the Line of Separation (LoS) the line where RK2 starts outperforming Euler. We Implemented a linear regression of the points where RK2 “crosses” E using as explanatory variable the input amplitude (Figure 6). It is noteworthy the separation between the region where “Euler wins” and the region where “RK2 wins” is consistent across model parameter sets. Figure 6 can then be used as a practical tool to assess, given an estimate of the FLOPS available and the range of possible inputs to the model, if known, which solver to use. This is an important result in neuromorphic engineering. In fact, the area under the Line of Separation is the most used for neuro-engineering purposes, for example in micro-controller applications. These results mean that, counterintuitively, it is always better for the parameters we explored to use E (in the FLOPS-input amplitude semi plane below the LoS) or RK2 (above the LoS) with a smaller discretization step than RK4 with a higher one. These results, that simpler methods (Euler and Runge Kutta 2) outperform more complex ones, is explained because the Izhikevich model presents a reset rule. The time stamp when this reset happens is considered the instant of the spiking event. Therefore, using a more precise solver with a larger discretization step, makes a bigger error in estimating the reset time than using a less precise one with a smaller discretization step. With higher FLOPS, instead, RK2’s is small enough to realize the best compromise between accuracy of the simulation of the membrane potential and time stepping for the reset rule. To check the robustness of the results, we used another quality metric, the Van Rossum Distance (see Fig. 7). This analysis yielded the same results as the VPD. We show the results for a specific Victor Purpura q parameter and Van Rossum t parameter, which are respectively q = 0.001/ms and t = 750 ms. Nonetheless, in the Appendix we show the same plots for different values of q and t, showing that out findings are robust to the selected cost.

**Figure 5:**
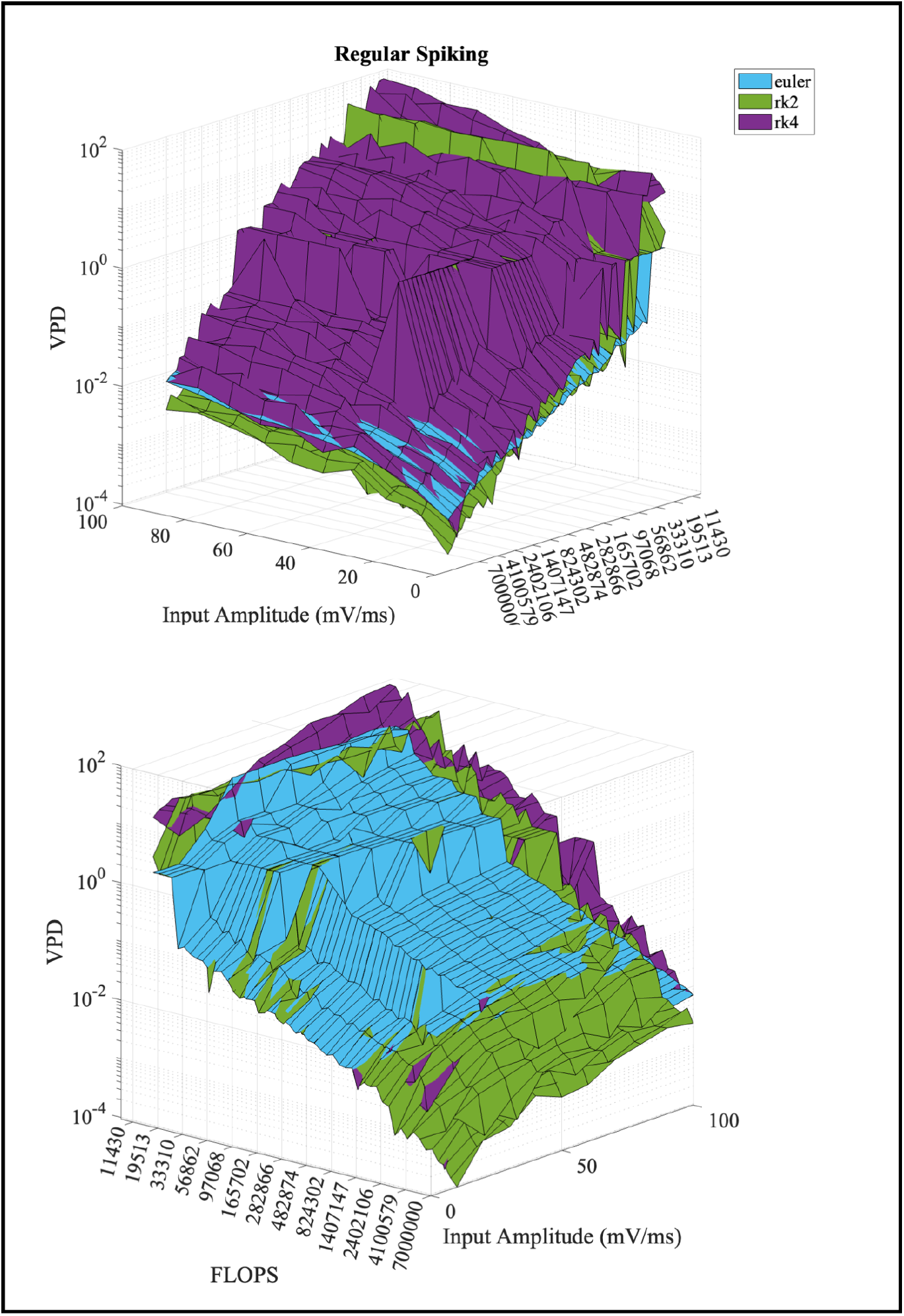
Each surface, coded by the color of the respective solver, represents the VPD from the reference as a function of the number of FLOPS and the Input Amplitude. The figures on the right show that RK4 is almost everywhere the solver with worst performance. On the left, we can see that there is a first blue area where Euler outperforms the other solvers, and a second green area where RK2 does. To assess quantitatively this area in all the 4 neuron models, see Figure 6 and Figure 7.

**Figure 6:**
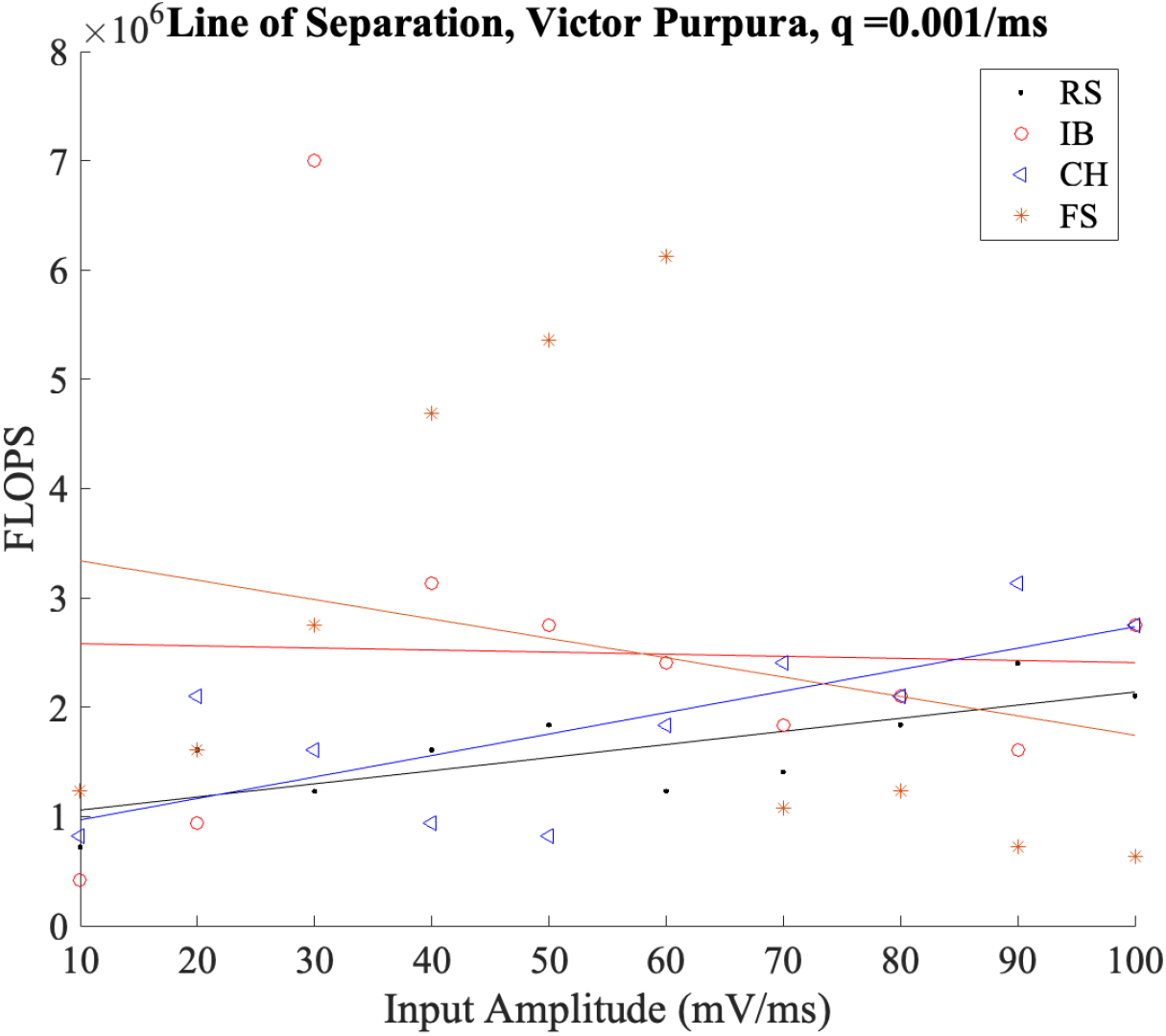
We implemented a linear fitting of the points where RK2 start outperforming Euler. Each color represents a different parameters set, following the same color coding of previous figures.

**Figure 7:**
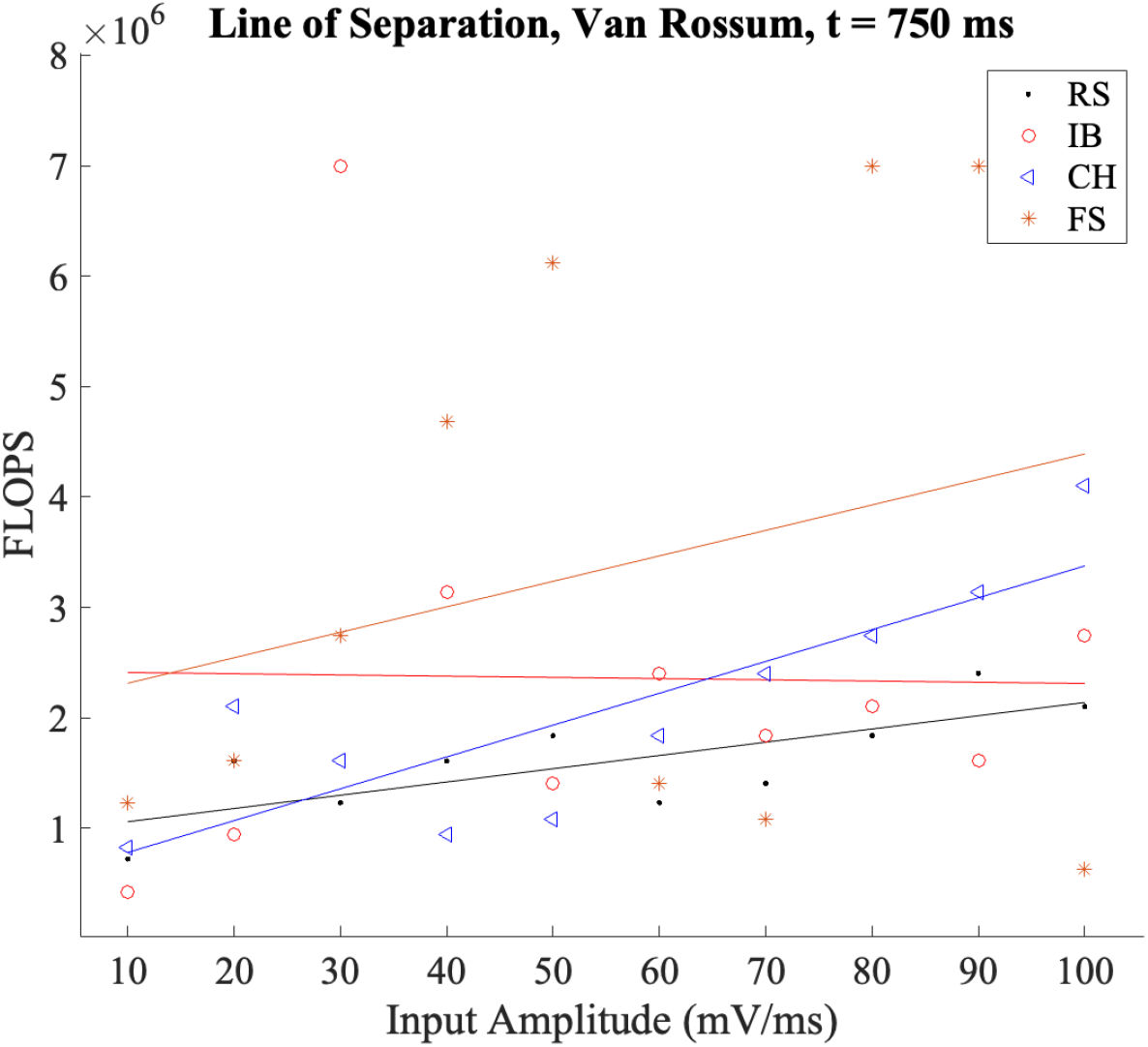
The same results of Figure 6 using the VRD instead of the VPD

This result was provided by considering as input to the model a step of current, as shown in Figure1, Figure 2. Being the model highly nonlinear, however, to assess whether this result is generalizable we tried as input sine functions of increasing frequency. Thus, keeping a fixed maximum amplitude of 30 mV/ms, we sweeped on the frequency of the sinusoid inputted to the model. The results in Figure 8 show that, even considering a broad range of frequencies, the primacy Euler-Runge Kutta 2 holds.

**Figure 8:**
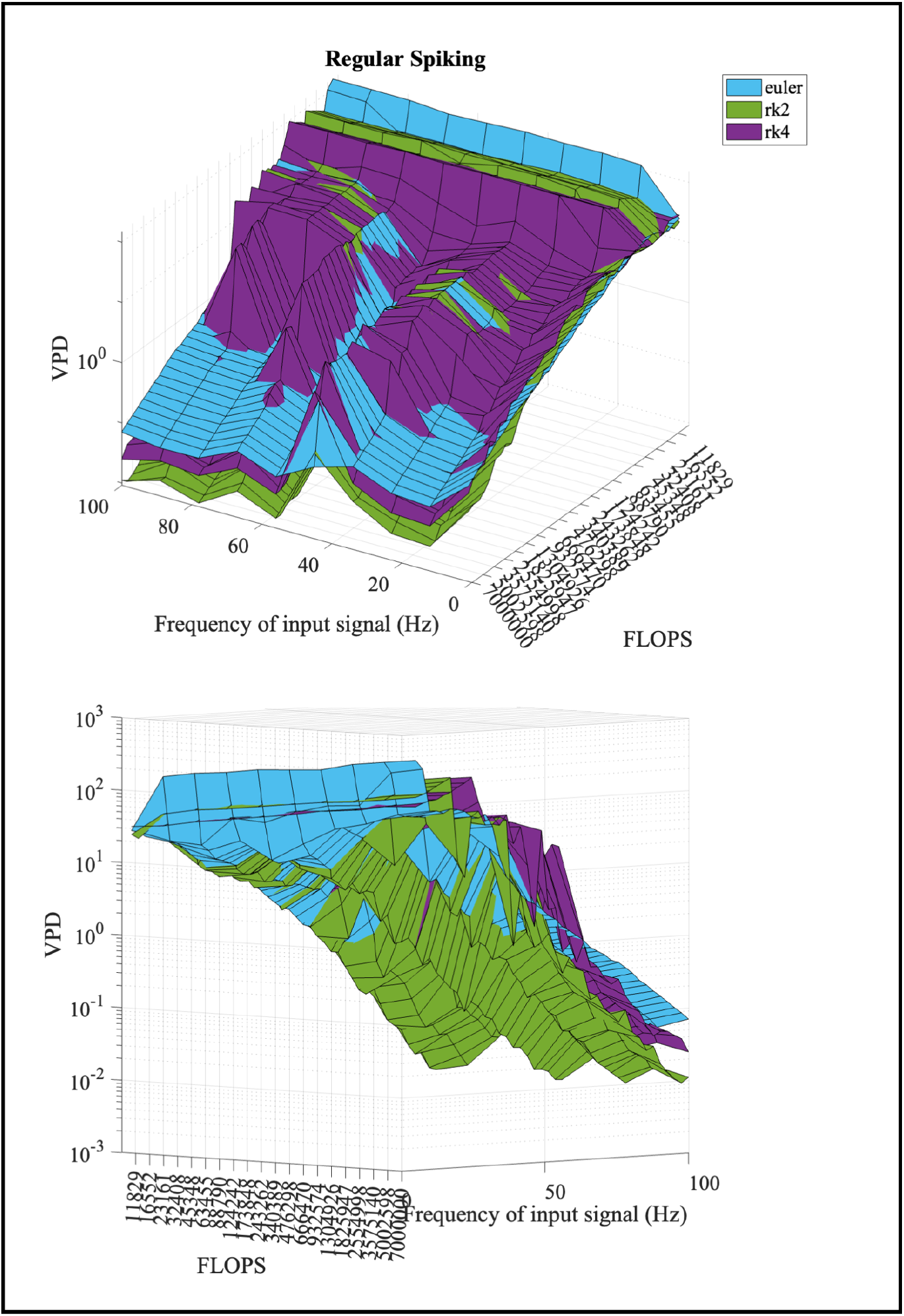
The same results of Figure 5. This time, however, we did not sweep on the input amplitude but on the frequency of the input sine function.

### 3.3 Reset Time

Finally, we quantified the duration of the OFF period necessary to reset the delay, as in Figure 4. To do so, we considered the squared Euclidean distance between the trajectory in the phase space of the numerical and the benchmark solutions. The time to achieve convergence was set as the time step when the distance was lower than a threshold of 1e-03 (see Figure 9). We repeated this analysis varying the input amplitude from 10 to 100 mV/ms. Results are shown for the maximum input amplitude since it provokes the highest firing rate and therefore the highest numerical instability. Moreover, we considered a range of 0.1-1 ms. Finally, wondering if the result was dependent on the duration of the spiking train, we ran the simulation with spike trains of different durations (from 100 ms to 1000 ms). It is noteworthy that the time for the solution to converge (the distance between the red line and the black line) does not depend on the duration of the spike train. Consider Figure 10. This result shows that the exponential decay of the distance is almost independent from the discretization time and the duration of the input. Both these last two images confirm that the numerical solution approaches a numerical limit cycle (thus accumulating the almost the same amount of delay for each spike, coherently with the previous results about the linearity) which takes almost the same time to converge to the numerical equilibrium solution.

**Figure 9:**
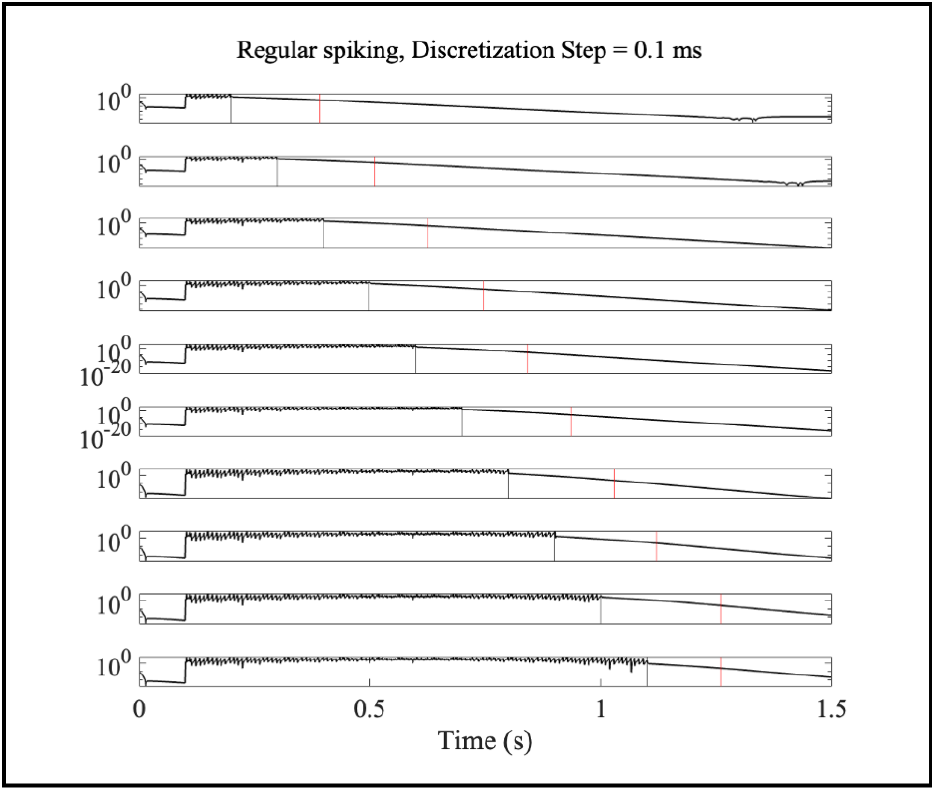
In the plots the discretization time is fixed, and each subplot represents a different duration of the spike train. The black vertical line is the end of the spike train, which coincides with the time step when the solution starts to converge. The red line is the time step when the distance becomes lower than a threshold, set to 0.001. The plots are logarithmic.

**Figure 10:**
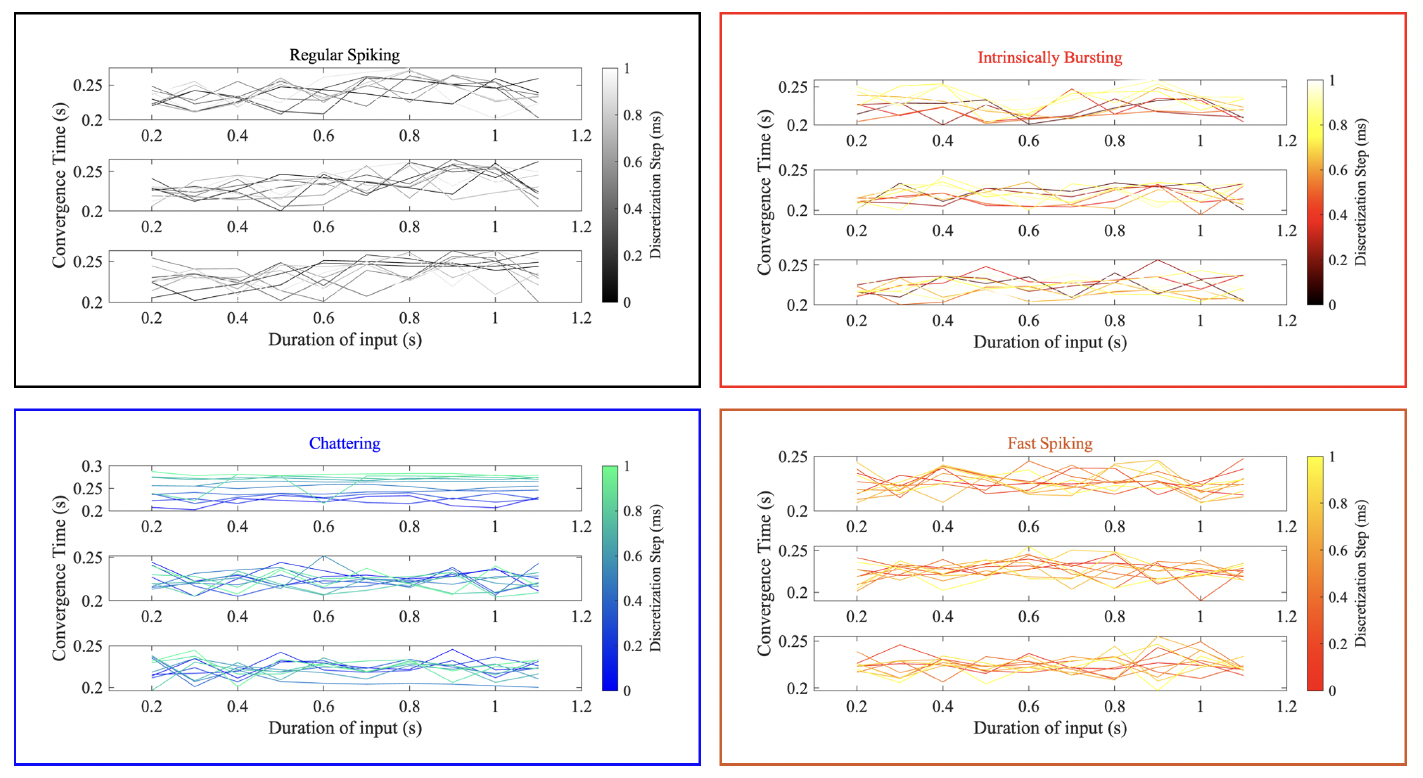
The convergence time, computed as described in Figure 9, for the three different ODE solvers, and by varying the input duration (therefore the duration of the spike train), in the x axis, and the discretization step (coded by the color).

## 4 Discussion and Conclusion

What makes the ISNM challenging from a mathematical perspective is the presence of a reset rule, which coincides with the spike. This rule made us question whether it was better to use a simpler solver with a lower time step, which allows to approximate the time of reset more accurately, or a more complex solver with a larger one. First, coherently with previous paper, we have shown the consequences of discretization on error accumulation. Secondly, we have proposed a simple pipeline to address this issue in a neuromorphic computing scenario. Being the major problem the reset rule and then the spike timing, we adopted spike train distances such as the Victor Purpura Distance. Nonetheless, this strategy can be extended with ease to different models (Brette and Gerstner [2005]) and different solvers, such as the Zero-Order Hold solver proposed by (Humphries and Gurney [2007]). Being spiking neuron models highly non-linear and strongly input dependent, we believe that this fast and easily reproducible approach, can be an effective and rapid method to assess the right solver for a neuro-engineering or computational neuroscience application. The primacy of simpler methods (Euler and RK2) is an encouraging result especially in micro-controller-based neuro-robotic applications. In fact, it allows for a greater grasp of all the operations being performed and is amenable of an extremely simple and mathematically understandable implementation. Moreover, we showed a way to analyze the time of convergence of the solver. It is useful to understand the time needed to achieve convergence and return to the equilibrium solution, and then, reset the error that gets accumulated during the spike train. We have found that this time is stable across different parameters, firing rates and solvers. Further developments of the work could conceive the generalization of the results to other models and the exploration of different solvers. We focused on fixed step solvers because they represent a simple and straightforward way to implement the model in micro-controllers and in simulations of large networks. However, these results can be extended with ease also to variable-step solvers.

## A Additional Figures

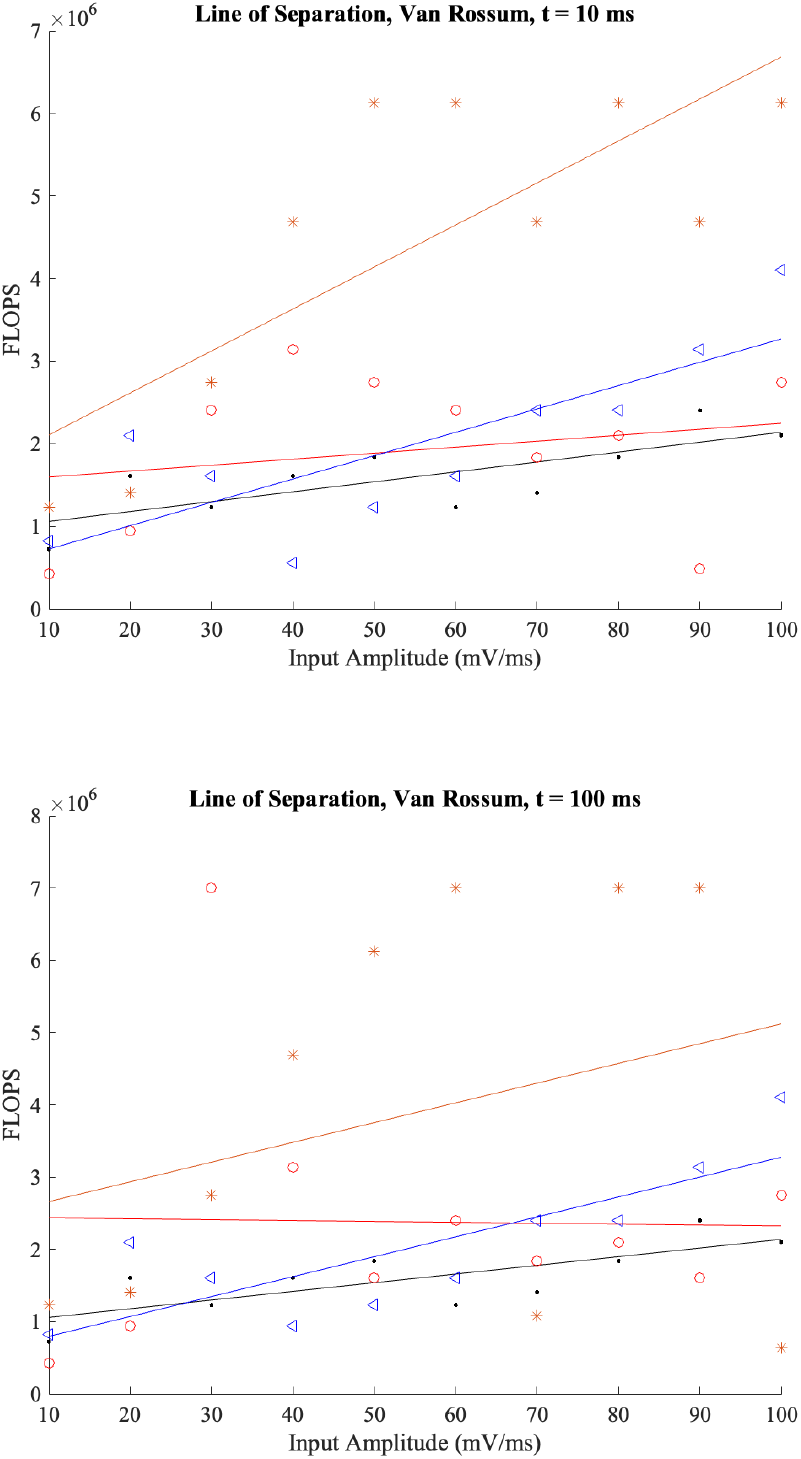

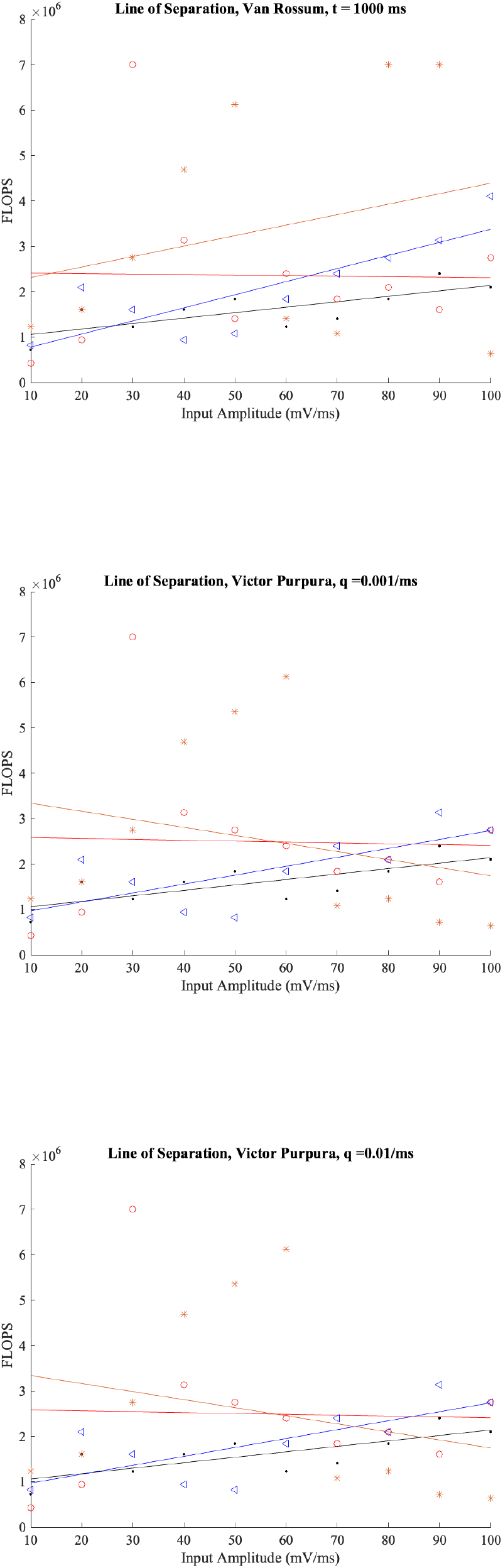

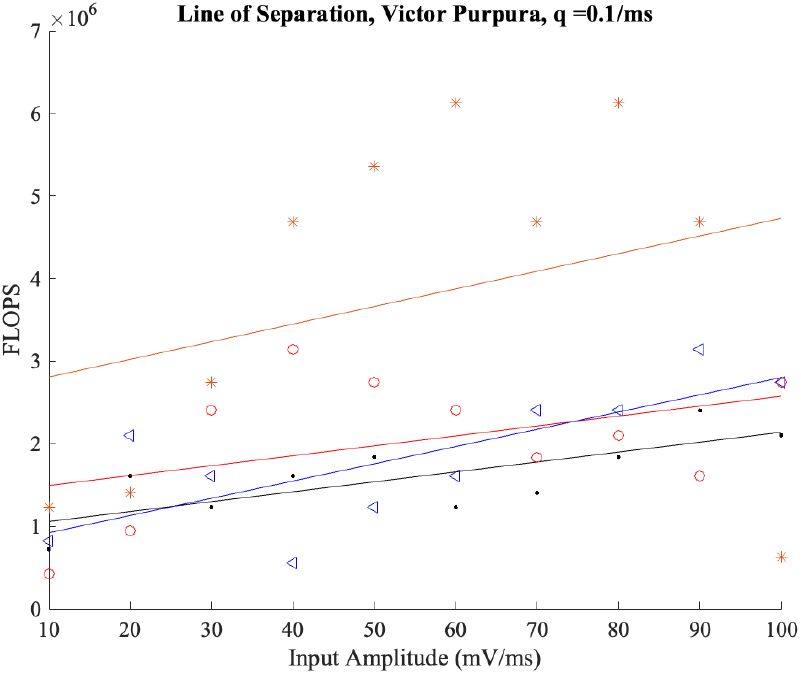

## References

Eugene M. Izhikevich. Simple model of spiking neurons, 2003. ISSN 10459227.

Eugene M. Izhikevich. Which model to use for cortical spiking neurons? IEEE Transactions on Neural Networks, 2004. ISSN 10459227. doi:10.1109/TNN.2004.832719.

Calogero Maria Oddo, Stanisa Raspopovic, Fiorenzo Artoni, Alberto Mazzoni, Giacomo Spigler, Francesco Petrini, Federica Giambattistelli, Fabrizio Vecchio, Francesca Miraglia, Loredana Zollo, Giovanni Di Pino, Domenico Camboni, Maria Chiara Carrozza, Eugenio Guglielmelli, Paolo Maria Rossini, Ugo Faraguna, and Silvestro Micera. Intraneural stimulation elicits discrimination of textural features by artificial fingertip in intact and amputee humans. eLife, 2016. ISSN 2050084X. doi:10.7554/eLife.09148.

Luke E. Osborn, Andrei Dragomir, Joseph L. Betthauser, Christopher L. Hunt, Harrison H. Nguyen, Rahul R. Kaliki, and Nitish V. Thakor. Prosthesis with neuromorphic multilayered e-dermis perceives touch and pain. Science Robotics, 2018. ISSN 24709476. doi:10.1126/scirobotics.aat3818.

Alberto Mazzoni, Calogero M. Oddo, Giacomo Valle, Domenico Camboni, Ivo Strauss, Massimo Barbaro, Gianluca Barabino, Roberto Puddu, Caterina Carboni, Lorenzo Bisoni, Jacopo Carpaneto, Fabrizio Vecchio, Francesco M. Petrini, Simone Romeni, Tamas Czimmermann, Luca Massari, Riccardo di Iorio, Francesca Miraglia, Giuseppe Granata, Danilo Pani, Thomas Stieglitz, Luigi Raffo, Paolo M. Rossini, and Silvestro Micera. Morphological Neural Computation Restores Discrimination of Naturalistic Textures in Trans-radial Amputees. Sci. Rep., 2020. ISSN 20452322. doi:10.1038/s41598-020-57454-4.

Xiumin Li, Hui Liu, Fangzheng Xue, Hongjun Zhou, and Yongduan Song. Liquid computing of spiking neural network with multi-clustered and active-neuron-dominant structure. Neurocomputing, 2017. ISSN 18728286. doi:10.1016/j.neucom.2017.03.022.

Kenneth L. Rice, Mohammad A. Bhuiyan, Tarek M. Taha, Christopher N. Vutsinas, and Melissa C. Smith. Fpga implementation of izhikevich spiking neural networks for character recognition. 2009. ISBN 9780769539171. doi:10.1109/ReConFig.2009.77.

Udaya Bhaskar Rongala, Alberto Mazzoni, and Calogero Maria Oddo. Neuromorphic artificial touch for categorization of naturalistic textures. IEEE Transactions on Neural Networks and Learning Systems, 28:819–829, 4 2017. ISSN 21622388. doi:10.1109/TNNLS.2015.2472477.

Mohadeseh Shafiei, Fatemeh Parastesh, Mahdi Jalili, Sajad Jafari, Matjaž Perc, and Mitja Slavinec. Effects of partial time delays on synchronization patterns in izhikevich neuronal networks. European Physical Journal B, 2019. ISSN 14346036. doi:10.1140/epjb/e2018-90638-x.

Mary Vinaya and Rose P. Ignatius. Effect of lévy noise on the networks of izhikevich neurons. Nonlinear Dynamics, 2018. ISSN 1573269X. doi:10.1007/s11071-018-4414-8.

Moslem Heidarpur, Arash Ahmadi, and Majid Ahmadi. Time step impact on performance and accuracy of izhikevich neuron: Software simulation and hardware implementation. 2020. doi:10.1109/iscas45731.2020.9180632.

Haitao Yu, Roberto F. Galán, Jiang Wang, Yibin Cao, and Jing Liu. Stochastic resonance, coherence resonance, and spike timing reliability of hodgkin–huxley neurons with ion-channel noise. Physica A: Statistical Mechanics and its Applications, 471:263–275, 4 2017a. ISSN 0378-4371. doi:10.1016/J.PHYSA.2016.12.039.

Haitao Yu, Rishi R. Dhingra, Thomas E. Dick, and Roberto F. Galán. Effects of ion channel noise on neural circuits: an application to the respiratory pattern generator to investigate breathing variability. Journal of neurophysiology, 117: 230–242, 1 2017b. ISSN 1522-1598. doi:10.1152/JN.00416.2016. URL https://pubmed.ncbi.nlm.nih.gov/27760817/.

Harish Gunasekaran, Giacomo Spigler, Alberto Mazzoni, Enrico Cataldo, and Calogero Maria Oddo. Convergence of regular spiking and intrinsically bursting izhikevich neuron models as a function of discretization time with euler method. Neurocomputing, 2019. ISSN 18728286. doi:10.1016/j.neucom.2019.03.021.

Giuseppe de Alteriis and Calogero Maria Oddo. Tradeoff between accuracy and computational cost of euler and runge kutta ode solvers for the izhikevich spiking neuron model. In 2021 10th International IEEE/EMBS Conference on Neural Engineering (NER), pages 730–733. IEEE, 2021.

Mark D. Humphries and Kevin Gurney. Solution methods for a new class of simple model neurons. Neural Computation, 2007. ISSN 08997667. doi:10.1162/neco.2007.19.12.3216.

Lyle N Long and Guoliang Fang. A review of biologically plausible neuron models. AIAA InfoTech@Aerospace Conference, 2010.

Michael J. Skocik and Lyle N. Long. On the capabilities and computational costs of neuron models. IEEE Transactions on Neural Networks and Learning Systems, 2014. ISSN 21622388. doi:10.1109/TNNLS.2013.2294016.

Michael Hopkins and Steve Furber. Accuracy and efficiency in fixed-point neural ode solvers. Neural Computation, 2015. ISSN 1530888X. doi:10.1162/NECO_a_00772.

Sergio Valadez-Godínez, Humberto Sossa, and Raúl Santiago-Montero. On the accuracy and computational cost of spiking neuron implementation. Neural Networks, 122:196–217, 2020.

Kuang-Hua Chang. Multiobjective optimization and advanced topics. e-Design, pages 1105–1173, 1 2015. doi:10.1016/B978-0-12-382038-9.00019-3.

Barry W. Connors and Michael J. Gutnick. Intrinsic firing patterns of diverse neocortical neurons. Trends in Neurosciences, 13:99–104, 3 1990. ISSN 0166-2236. doi:10.1016/0166-2236(90)90185-D.

Jay R. Gibson, Michael Belerlein, and Barry W. Connors. Two networks of electrically coupled inhibitory neurons in neocortex. Nature 1999 402:6757, 402:75–79, 11 1999. ISSN 1476-4687. doi:10.1038/47035. URL https://www.nature.com/articles/47035.

Jonathan D. Victor and Keith P. Purpura. Nature and precision of temporal coding in visual cortex: A metric- space analysis. Journal of Neurophysiology, 1996. ISSN 00223077. doi:10.1152/jn.1996.76.2.1310.

Eero Satuvuori and Thomas Kreuz. Which spike train distance is most suitable for distinguishing rate and temporal coding? Journal of Neuroscience Methods, 2018. ISSN 1872678X. doi:10.1016/j.jneumeth.2018.02.009.

Romain Brette and Wulfram Gerstner. Adaptive exponential integrate-and-fire model as an effective description of neuronal activity. Journal of Neurophysiology, 94:3637–3642, 11 2005. ISSN 00223077. doi:10.1152/JN.00686.2005/ASSET/IMAGES/LARGE/Z9K0110549990003.JPEG. URL https://journals.physiology.org/doi/abs/10.1152/jn.00686.2005.

